# Direct quantification of chemogenetic H_2_O_2_ production in live human cells

**DOI:** 10.1101/2023.05.03.539306

**Authors:** Wytze T. F. den Toom, Daan M. K. van Soest, Paulien E. Polderman, Miranda H. van Triest, Lucas J. M. Bruurs, Sasha De Henau, Boudewijn M. T. Burgering, Tobias B. Dansen

**Affiliations:** Center for Molecular Medicine, University Medical Center Utrecht, Universiteitsweg 100, 3584 CG Utrecht, The Netherlands; Hubrecht Institute, Uppsalalaan 8,3584 CT Utrecht, The Netherlands; Oncode Institute, Jaarbeursplein 6, 3521 AL Utrecht, The Netherlands

## Abstract

Reactive Oxygen Species (ROS) in the form of H_2_O_2_ can act both as physiological signaling molecules as well as damaging agents, depending on its concentration and localization. The downstream biological effects of H_2_O_2_ were often studied making use of exogenously added H_2_O_2_, generally as a bolus and at supraphysiological levels. But this does not mimic the continuous, low levels of intracellular H_2_O_2_ production by for instance mitochondrial respiration. The enzyme D-Amino Acid Oxidase (DAAO) catalyzes H_2_O_2_ formation using D-amino acids, which are absent from culture media, as a substrate. Ectopic expression of DAAO has recently been used in several studies to produce inducible and titratable intracellular H_2_O_2_. However, a method to directly quantify the amount of H_2_O_2_ produced by DAAO has been lacking, making it difficult to assess whether observed phenotypes are the result of physiological or artificially high levels of H_2_O_2_. Here we describe a simple assay to directly quantify DAAO activity by measuring the oxygen consumed during H_2_O_2_ production. The oxygen consumption rate of DAAO can directly be compared to the basal mitochondrial respiration in the same assay, allowing to estimate whether the ensuing level of H_2_O_2_ production is within the range of physiological mitochondrial ROS production. We show that the assay can also be used to select clones that express differently localized DAAO with the same absolute level of H_2_O_2_ production to be able to discriminate the effects of H_2_O_2_ production at different subcellular locations from differences in total oxidative burden. This method therefore greatly improves the interpretation and applicability of DAAO-based models, thereby moving the redox biology field forward.

## Introduction

H_2_O_2_ is produced endogenously at several subcellular locations, either directly or indirectly as the more reactive superoxide anion (O_2_^•-^) that is subsequently dismutated to form H_2_O_2_. The main intracellular sources of H_2_O_2_ are the mitochondrial electron transport chain (ETC) and NADPH oxidases located in the plasma membrane, endoplasmic reticulum, and nucleus (Wong et al., 2019). Cell fate downstream of redox signaling may range from enhanced proliferation (Kirova et al., 2022) and differentiation (De Henau et al., 2020; Sato et al., 2014; Schader et al., 2020) to induction of cell death dependent on where and how much H_2_O_2_ is produced (Bleier et al., 2015; Cheung et al., 2016), reviewed in (Sies et al., 2022). The Redox Signaling field has rapidly emerged following the discovery that endogenously formed H_2_O_2_ serves as a second messenger. Redox signaling proceeds through the specific and reversible oxidation of dedicated cysteines in proteins, and over the past few decades the involvement of redox signaling downstream of H_2_O_2_ in the regulation of a wide range of vital processes has become apparent (Holmstrom and Finkel, 2014; Sies and Jones, 2020; Winterbourn, 2018).

With the development of sensitive techniques to detect endogenous H_2_O_2_ (Bilan and Belousov, 2018; Morgan et al., 2016; Pak et al., 2020; Schwarzlander et al., 2016), it became clear that endogenous H_2_O_2_ is characterized by steep concentration gradients, and hence does not diffuse far from its site of production (De Henau et al., 2020; Ermakova et al., 2019; Hoehne et al., 2022; Mishina et al., 2019; Pak et al., 2020; Winterbourn, 2018). These observations also strengthened the notion that redox signaling should preferably be studied under conditions that best mimic physiological circumstances (Murphy et al., 2022). Hence, using localized, low levels of steady state H_2_O_2_ production rather than bolus addition of exogenous H_2_O_2_. To this end the redox biology field has embraced a chemogenetic approach to achieve inducible, titratable, and targetable endogenous H_2_O_2_ production that makes use of the ectopic expression of the enzyme D-amino acid oxidase (DAAO) from the yeast *R. gracilis* (Haskew-Layton et al., 2010; Matlashov et al., 2014; Stegman et al., 1998). Administration of D-amino acids, which are normally (largely) absent from culture media and animal model systems, is subsequently used to start H_2_O_2_ production (Figure 1). The DAAO system has been used recently in a variety of studies aimed at e.g., measuring whether H_2_O_2_ produced in the mitochondrial matrix leaks into the cytoplasm or to study the isolated effect of oxidative damage in the hearts of rats in the absence of other cardiac pathology. A (non-exhaustive) overview of recent studies applying DAAO to generate H_2_O_2_ in vivo or in live cells is provided in table 1.

**Table 1:** A selection of recent studies using ectopic expression of *R. gracilis* DAAO.

| Study topic | References |
| --- | --- |
| Astrocyte-dependent H<sub>2</sub>O<sub>2</sub>-induced neuroprotection | (Sies, 2021) |
| DAAO as putative anti-cancer therapy | (Fang et al., 2002)<br>(Fuentes-Baile et al., 2020)<br>(Li et al., 2008)<br>(Stegman et al., 1998) |
| H<sub>2</sub>O<sub>2</sub> transport by Aquaporin 8 | (Kruger et al., 2021) |
| Mitochondrial H<sub>2</sub>O<sub>2</sub> dynamics and release | (Kruger et al., 2021)<br>(Moon et al., 2020)<br>(Pak et al., 2020)<br>(Stein et al., 2020) |
| Oxidative heart failure in rats and mice | (Nanadikar et al., 2023)<br>(Sorrentino et al., 2019)<br>(Spyropoulos et al., 2022)<br>(Steinhorn et al., 2018) |
| Role of H<sub>2</sub>O<sub>2</sub> gradients in zebrafish neuronal development and neuronal health | (Amblard et al., 2020)<br>(Haskew-Layton et al., 2010) |
| ROS-dependent cell cycle regulation | (Kirova et al., 2022) |

**Figure 1:**
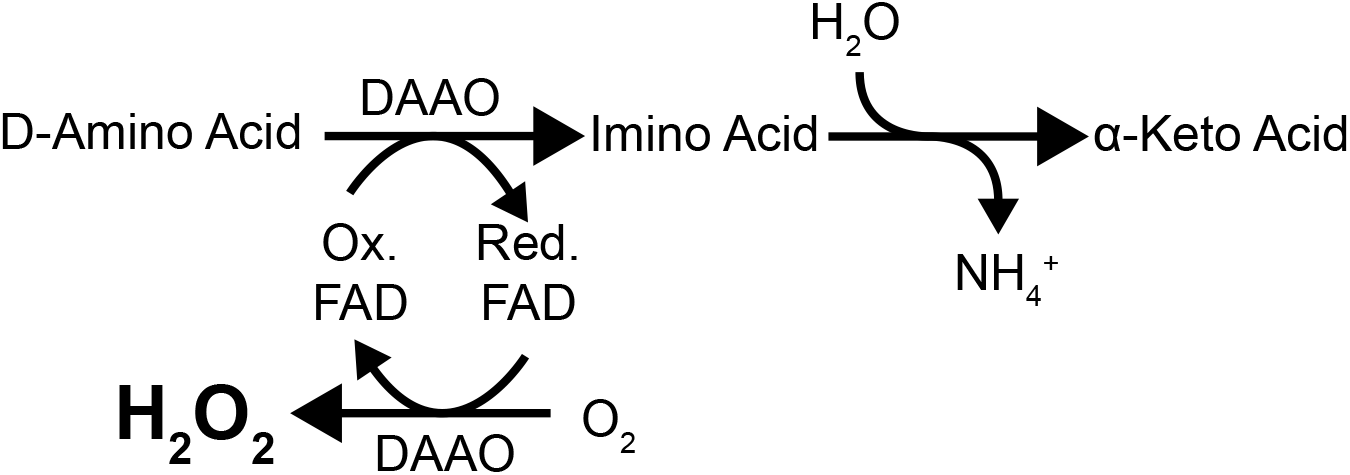
Schematic of the reaction catalyzed by DAAO. DAAO converts D-amino acids to imino acids, reducing its cofactor FAD. FAD is subsequently oxidized, generating H_2_O_2_ while consuming equimolar amounts of O_2_. The formed imino acids spontaneously hydrolyze to α-keto acids.

Many of these studies make use of fluorescence-based H_2_O_2_ sensors like HyPer7 as a means to quantify H_2_O_2_ production by DAAO, either expressed as a fusion protein of DAAO or expressed separately, for instance with a different localization tag to assess H_2_O_2_ gradients across organelles (Erdogan et al., 2021; Matlashov et al., 2014; Saeedi Saravi et al., 2020). Although this approach can report on relative H_2_O_2_ levels within one model system, it may not be suitable to directly compare DAAO activity, and therefore H_2_O_2_ levels, between for instance different cell lines or when DAAO is expressed at different subcellular sites. DAAO activity is dependent on its expression level, localization, presence of the cofactor FAD and availability of its substrate D-amino acids. The latter will likely not only depend on how much substrate has been administered to the culture media, but also on its uptake, and diffusion and transport rates across organellar membranes. Furthermore, genetically encoded fluorescent sensors like HyPer7 report on the *combined* rates of oxidation (by H_2_O_2_) and reduction (by the Trx system in case of HyPer7) (Kritsiligkou et al., 2021), and therefore its ratiometric readout depends not only on DAAO activity but also on the local reductive capacity, which varies between organelles and cell lines (Sies, 2021). Hence, absolute levels of H_2_O_2_ produced by DAAO and H_2_O_2_ concentration measured with for instance HyPer7 may not necessarily be directly proportional, and this may influence conclusions drawn. Besides, HyPer7 itself could in principle affect the reductive capacity by acting as an H_2_O_2_ sink, making comparisons between conditions and cell lines even more difficult.

We have developed an assay that reports directly on DAAO activity independent of the local reductive capacity. Since H_2_O_2_ production by DAAO consumes an equimolar amount of oxygen, DAAO activity can be determined by measuring the oxygen consumption of cells upon the addition of D-amino acids using for instance a Seahorse XFe Analyzer, which is a widely available piece of equipment in research institutes. We show that this method can be used to quantify DAAO activity in monoclonal RPE1-hTERT cell lines that express the enzyme at various subcellular sites. This allowed us to determine and directly compare at what levels H_2_O_2_ impairs viability when produced in different compartments. We show that intramitochondrial H_2_O_2_ production prevents outgrowth of cells at much lower levels as compared to production at the outer mitochondrial membrane or near the genomic DNA. We also provide evidence that the uptake of D-amino acids competes with that of L-amino acids and that D-amino acid uptake is a rate-limiting step in intracellular DAAO-dependent H_2_O_2_ production. We think that our OCR-based approach to directly quantify H_2_O_2_ production levels is of value to the redox biology community and could be applied to the quantification of other enzyme activities that consume or produce oxygen as a byproduct.

## Results

### Oxygen consumption rate can be used as a proxy for DAAO activity

We hypothesized that if the activity of ectopically expressed DAAO would be high enough as compared to other oxygen dependent cellular reactions, DAAO activity could be directly monitored by measuring the oxygen consumption rate (OCR) in a Seahorse XFe24 analyzer. The standard injection ports of the machine can be used for the addition of D-amino acids, allowing for baseline and induced OCR measurements within the same run. We used D-Ala with the rationale that its other reaction products besides H_2_O_2_ (i.e. pyruvate and ammonia) are abundantly present and rapidly metabolized, and therefore relatively minor increases in the flux of these metabolites would have negligible effects on cell physiology or OCR. Most previous studies using DAAO also used D-Ala as a substrate. To minimize variation, in this study we used monoclonal Human RPE1-hTert cells that stably expressed *R. gracilis* DAAO (without its C-terminal peroxisomal localization signal SKL) fused to various localization tags as well as mScarlet. Localization tags used in this study can be found in table 2.

**Table 2:** Utilized localization tags to target DAAO.

| abbreviation | Localization tag and targeted compartment | Model for which endogenous $H_2O_2$ source |
| --- | --- | --- |
| TOM20 | Targeting sequence of human TOMM20; cytoplasmic side of the outer mitochondrial membrane | Release of $H_2O_2$ from mitochondria |
| H2B | Human Histone 2B; Nucleosome | $H_2O_2$ production close to the genomic DNA (e.g. as done by LSD1) |
| MLS | 2 x Presequence of human COX-8; Mitochondrial matrix | $H_2O_2$ production in the the mitochondrial matrix. |
| IMS | Presequence of human DIABLO; Mitochondrial Intermembrane space | $H_2O_2$ production in the mitochondrial intermembrane space. |

Figure 2A shows that an increase in OCR upon addition of D-Ala (using L-Ala as a control) can indeed be observed in RPE1-hTert^TOM20-DAAO^ cells. At 20 mM D-Ala, the H_2_O_2_ production is about 60 pmol/min which translates to ∼1.5 fmol cell^-1^ min^-1^. OCR before addition of D-Ala consists predominantly of mitochondrial respiration as this can be largely inhibited by the ATP synthase inhibitor Oligomycin (Figure 1A). Metabolic differences between wells are expected to be largely eliminated by addition of Oligomycin, yielding a more reliable baseline that correlates better with cell number. Furthermore, DAAO-induced OCR can in this way be compared to the OCR that stems from mitochondrial respiration, to estimate whether the observed rates are near-physiological or not. It has been estimated that under basal conditions about 1% of oxygen passing through the ETC ends up as superoxide (Chance et al., 1979; Murphy, 2009), and two superoxide molecules will generate one molecule of H_2_O_2_ (catalyzed by SOD). Figure 2B shows that oligomycin treatment indeed provides a larger relative increase above baseline, allowing for more robust measurements.

**Figure 2:**
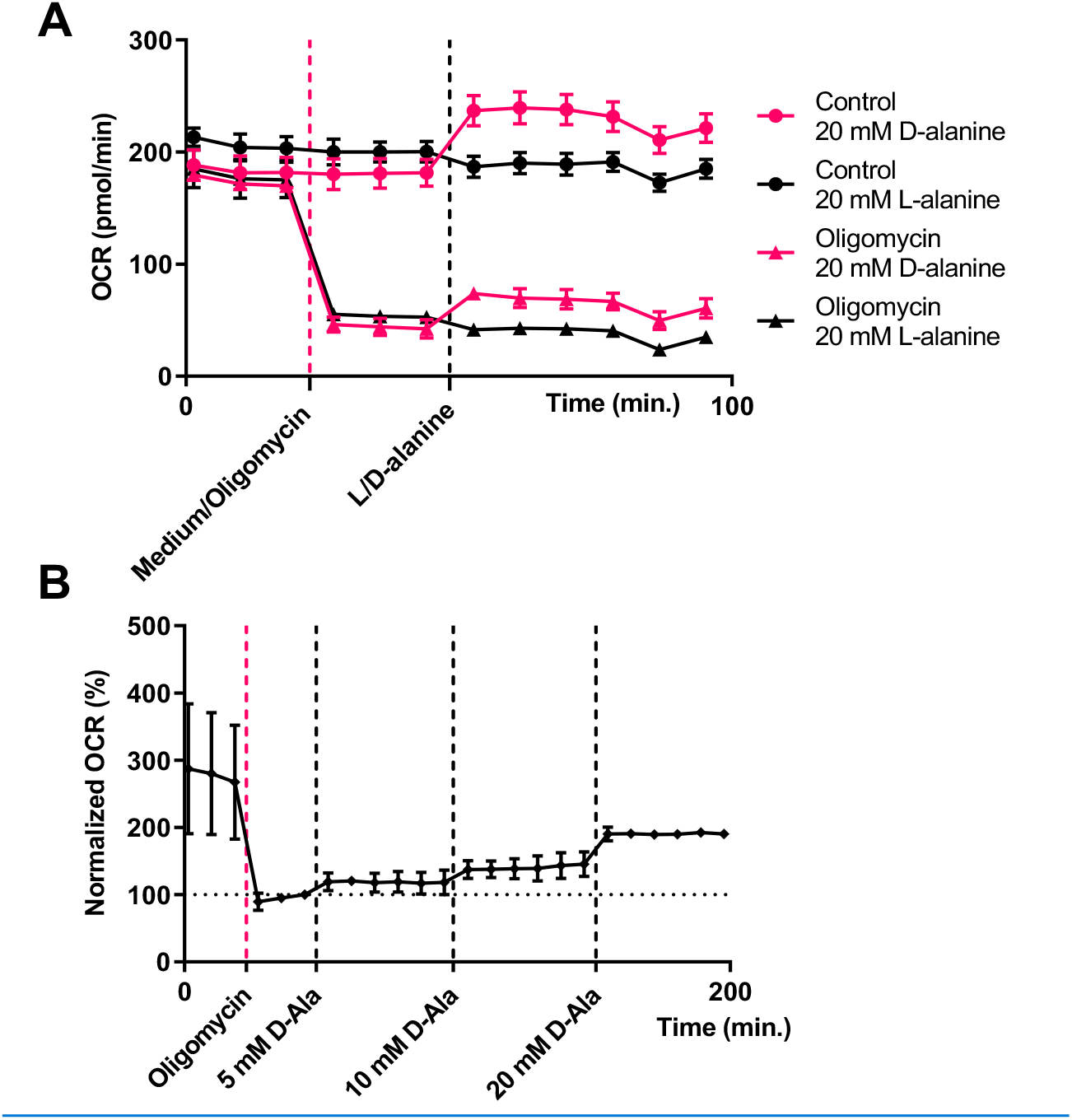
H_2_O_2_ production by ectopically expressed DAAO can be estimated by measuring oxygen consumption in a Seahorse XF analyzer. A. RPE1-hTert^TOM20-DAAO^ cells treated with L- or D-Ala in the presence and absence of oligomycin. Treatment with D-Ala, but not L-Ala leads to increased OCR. Oligomycin inhibits mitochondrial respiration, resulting in more reliable estimation of DAAO-dependent OCR with lower signal to noise. The error bars represent the standard deviation of 2 - 3 technical replicates (multiple wells per condition in one Seahorse plate). Typical result of several biological replicates. B. Sequential injections show [D-Ala] dependent increases in the mean OCR of RPE1-hTert^TOM20-DAAO^ cells. The third timepoint after oligomycin addition was set to 100%. Note that addition of oligomycin decreases the variability of the measurements between wells. The indicated [D-Ala] on the horizontal axis represents the total [D-Ala] in the well after each injection. Error bars: standard deviation of 5 technical replicates.

Each well in a Seahorse XF Analyzer has four injection ports. DAAO activity can thus be titrated using increasing D-amino acid concentrations in a single well. This allows for the comparison of the DAAO activity across a range of incremental D-alanine concentrations, without having to use different wells (and normalization) for each D-alanine concentration (Figure 2B), the oligomycin-normalized oxygen consumption of RPE1-hTert^TOM20-DAAO^ cells is shown at different concentrations of D-alanine. Figure 2B also shows that normalization to the OCR after oligomycin treatment eliminates most variation across different wells. The DAAO-dependent OCR is linearly proportional to the applied extracellular concentration of D-Ala over the tested range (0-20 mM). A plateau in OCR is observed for each concentration, suggesting that also at the lowest applied concentration substrate does not become limiting over the time-course of the measurement (50 minutes following each addition).

### The HyPer7 signal is dependent on glucose availability

Glucose metabolism is a major source of the reducing equivalent NADPH, and HyPer7 reports on the combined rate of oxidation (by H_2_O_2_) and reduction (by Trx). We compared the cytosolic NES-HyPer7 signal in response to H_2_O_2_ produced by DAAO^TOM20^ cultured in media containing either 1 g/L or 4.5 g/L glucose. Indeed, the oxidized/reduced HyPer7 ratio in low glucose media is much higher as compared to those in high glucose media at the same concentration of D-Ala (Figure 3). Because the rate of H_2_O_2_ production can be deduced from the OCR experiments, the relative reductive capacity of the cytoplasm under different conditions can be estimated from the HyPer7 ratio. Under low glucose conditions the HyPer7 ratio increases roughly twice as fast as under high glucose conditions at [D-Ala] ≤ 10 mM. Above 20 mM [D-Ala] or when a bolus of exogenous H_2_O_2_ is provided oxidation of the HyPer7 probe seems to outcompete reduction also under high glucose conditions. This example also illustrates that HyPer7 indeed reports on H_2_O_2_ dependent changes in the ratio of oxidation and reduction and not directly on H_2_O_2_ levels, and that care should be taken to keep culture conditions identical when HyPer7 is used to estimate or compare H_2_O_2_ levels.

**Figure 3:**
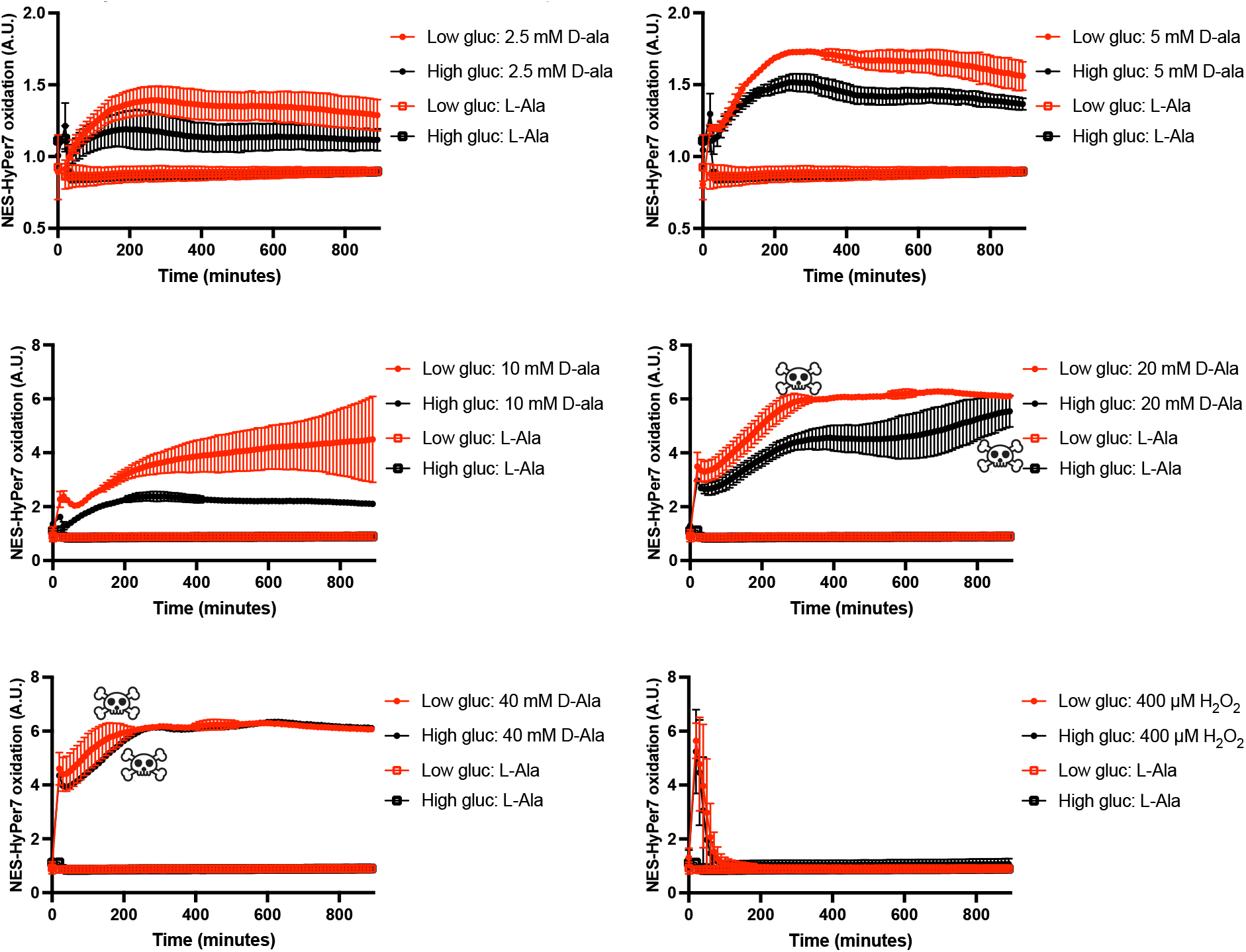
HyPer7 is not solely dependent on DAAO-activity but also glucose availability. Measurements of NES-HyPer7 oxidation in RPE1-hTERT-DAAO^TOM20^ cells cultured in either low glucose (1g/L) or high glucose (4.5g/l) conditions. The skull symbols indicate the induction of massive cell death based on cellular morphology. Upon activation of DAAO, HyPer7 oxidation increases more rapidly and to a higher level in low glucose medium compared to high glucose. Only at very high oxidative insults (>20 mM D-Ala or 400 μM of exogenous H2O2) is a similar oxidation of HyPer7 detected. This illustrates that HyPer7 signal can be influenced by other factors besides DAAO activity like glucose availability.

### Transmembrane transport of D-alanine is a rate limiting factor for DAAO activity

Based on the reported *in vitro* Michaelis-Menten constant for DAAO using D-Ala as a substrate (2.6 mM)(Rosini et al., 2009), one would expect that DAAO activity would be ∼75% saturated at 5 mM and almost 95% saturated at 10 mM of D-Ala. Our experiments in live cells however suggest a linear increase in DAAO activity upon an extra addition of 10 mM D-Ala (to yield a total of 20 mM) and hence an apparent K_M_ that is much higher than 2.6 mM. To test whether the rate of import of D-Ala over the plasma membrane could be a limiting factor for DAAO activity we permeabilized 20 mM D-Ala treated RPE1-hTert^TOM20-DAAO^ cells in the Seahorse machine by adding Saponin or Digitonin from the third injection port. Indeed, permeabilization leads to a rapid increase in DAAO-dependent OCR (Figure 4A). D-Ala likely uses the same amino acid importer as L-Ala, and several other amino acids (Hatanaka et al., 2002). Indeed, an injection with an excess of L-Ala subsequently decreased the D-Ala induced OCR, suggesting that L-Ala and D-Ala compete for the same import machinery (Figure 4B).

**Figure 4:**
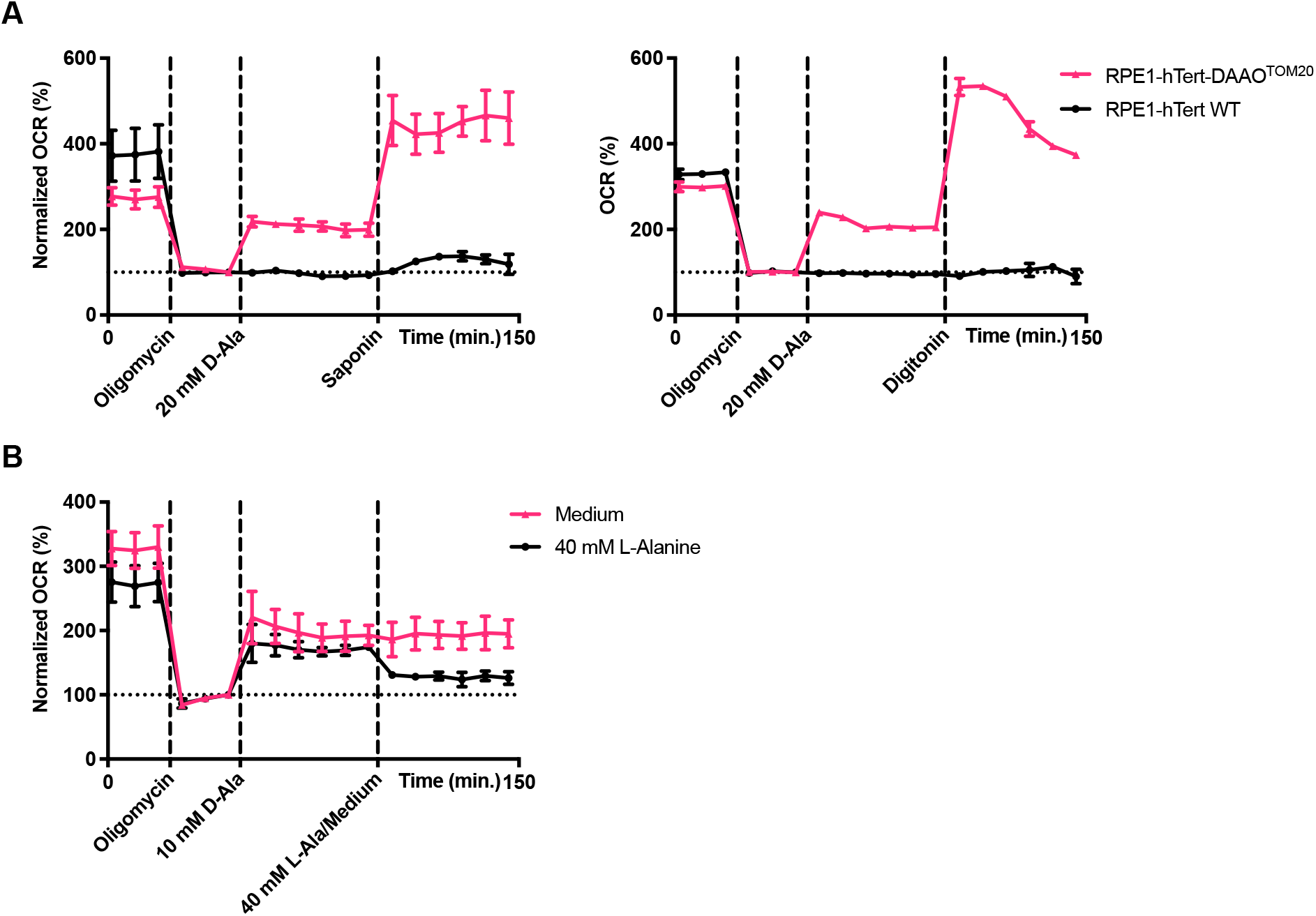
Transmembrane transport of D-Ala is rate limiting factor for DAAO activity A. Normalized OCR of RPE1-hTert^TOM20-DAAO^ and parental RPE1-hTert cells treated with oligomycin, 20 mM D-alanine and 25 μg/mL saponin or digitonin. These concentrations of saponin and digitonin in a Seahorse plate lead to permeabilization of the cell membrane(Salabei et al., 2014). Permeabilization leads to a rapid ∼two-fold increase in DAAO activity, indicating that transmembrane transport of D-alanine is a limiting factor in DAAO activity. The third timepoint after oligomycin addition was set to 100%. The error bars represent the standard deviation of 2 technical replicates. B. Normalized OCR of RPE1-hTert^TOM20-DAAO^ cells treated with oligomycin, 10 mM D-alanine and 40 mM L-Alanine (medium as control). The addition of a surplus of L-Ala decreases DAAO activity, suggesting L-Alanine competes with D-Alanine to be transported over the membrane. The third timepoint after oligomycin addition was set to 100%. The error bars represent the standard deviation of 2 technical replicates.

### OCR analysis enables the calibration of cell lines expressing DAAO in different compartments

We next used our Seahorse technology-based assay to calibrate and select monoclonal RPE1-hTert cell lines that have similar DAAO activity, but that express DAAO at different subcellular locations. These cell lines could help to elucidate whether an H_2_O_2_-induced phenotype is indeed the result of differential localized production or due to a difference in DAAO activity and ensuing overall oxidative burden. To this end we lentivirally transduced RPE1-hTert cells with constructs expressing DAAO fused to various localization tags (See Table 2) and generated monoclonal lines. Figure 5B shows that different mono-clonal lines expressing DAAO targeted to the same location can indeed have different DAAO activities.

**Figure 5:**
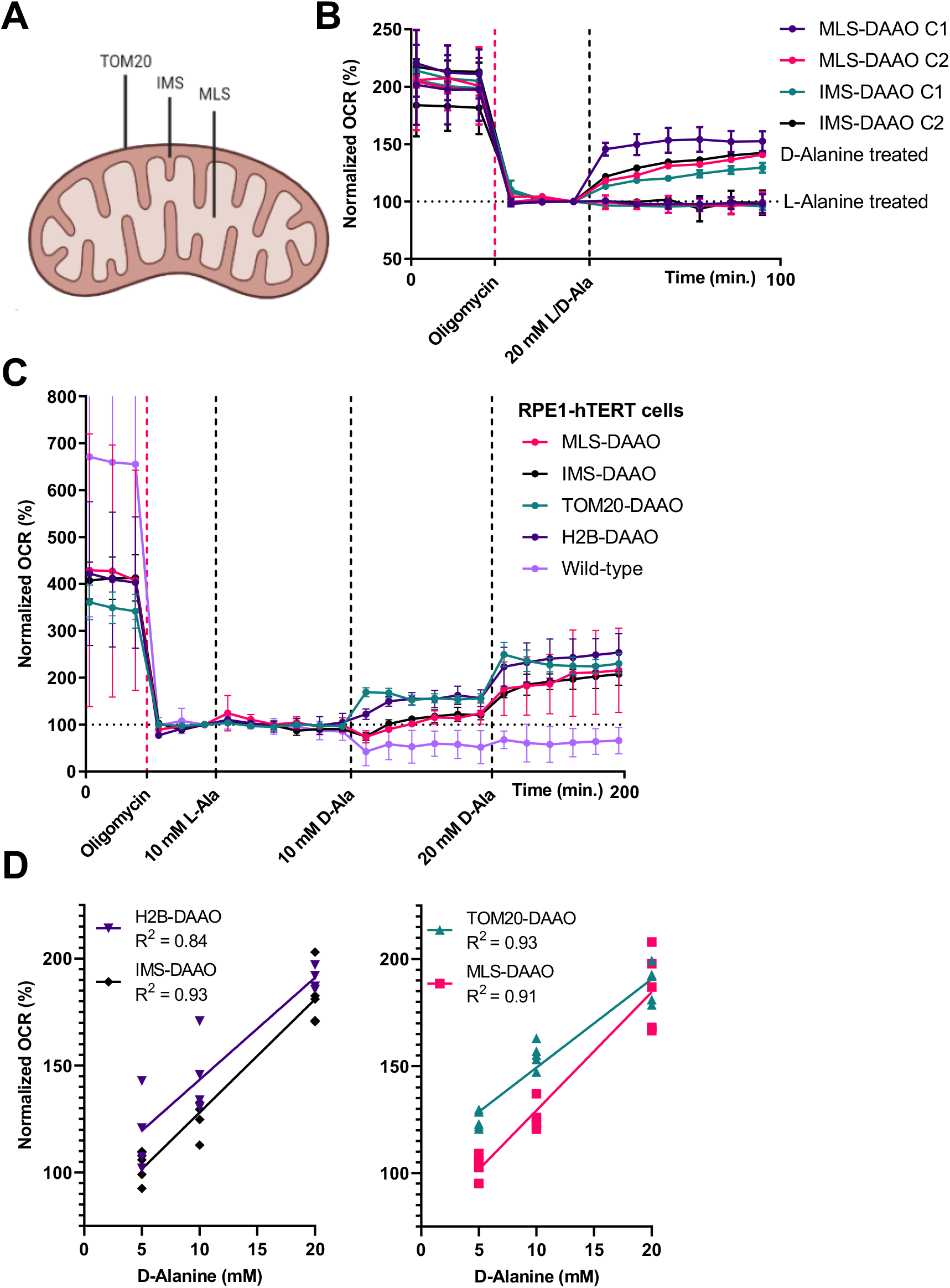
MLS-, IMS-, TOM20- and H2B-DAAO cells produce comparable levels of H_2_O_2_. A. Schematic overview of the localization of TOM20-, IMS- and MLS-DAAO constructs. H2B-DAAO is attached to nucleosomes in the nucleus (not shown). B. Normalized OCR of different MLS- and IMS-DAAO clones. MLS-DAAO clone 2 and IMS-DAAO clone 2 cells have a very similar level of oxygen consumption/DAAO activity at 20 mM D-alanine. The third timepoint after oligomycin addition was set to 100%. The error bars represent the standard deviation of 2 - 3 technical replicates.

Clones with similar DAAO activities were selected based on OCR measurements using increasing [D-Ala]. RPE1-hTert-DAAO^IMS^ clone 2 and RPE1-hTert-DAAO^MLS^ clone 2 had very similar levels of DAAO activity (Figure 5B), and so did the RPE1-hTert-DAAO^H2B^ and RPE1-hTert-DAAO^TOM20^ clones but note that the activity of the MLS and IMS localized DAAO was lower as compared to H2B and TOM20 localized DAAO (Figure 5C-D), and we did not find clones with equal activity at all four localizations. The RPE1-hTert-DAAO^H2B^ and RPE1-hTert-DAAO^TOM20^ cells consumed an amount of oxygen equivalent to roughly 20% of the basal mitochondrial respiration at 10 mM D-alanine. For MLS- and IMS-DAAO cells, this was 7%. These numbers are much higher than what can likely be achieved physiologically. Lowering [D-Ala] further will result in (near) physiological H_2_O_2_ levels, but these would be more difficult to pick up in this assay. The lower DAAO activity in the RPE1-hTert-DAAO^IMS^ and RPE1-hTert-DAAO^MLS^ lines should be considered when comparing the effects of DAAO-derived H_2_O_2_. Using a range of [D-Ala] in experiments could help to achieve similar rates of H_2_O_2_ production in all four lines.

### The site of H_2_O_2_ production determines its lethal dose

An advantage of using these OCR-based calibrated monoclonal RPE1-hTert-DAAO cell lines is the possibility to directly compare phenotypes downstream of equal but differentially localized H_2_O_2_ production. This may uncover for instance differences in local reductive capacity or the engagement of different signaling cascades. As an example, we have used the four monoclonal RPE1-hTert-DAAO cell lines to measure whether changes in cell density in response to localized H_2_O_2_ production depends on the site of production. We use ‘cell density’ because the Crystal Violet assay does not distinguish between differences in proliferative rate and cell death. Cell density was also somewhat decreased at higher levels of the L-Ala control in all four lines, which could be due to competition with essential amino acids for the same transporter. This observation suggests that care should be taken when studying the effects of redox signaling on cell cycle using the DAAO system, and that ideally equimolar amounts of L-Ala should be used as a control for D-Ala. Production of H_2_O_2_ at the nucleosome or the cytoplasmic side of the outer mitochondrial membrane shows a sudden decrease in cell density at ∼20 mM D-Ala, whereas H_2_O_2_ production in the mitochondrial intermembrane space or matrix showed a more gradual drop in viability, starting already at 2.5 mM D-Ala (Figure 5 & Sup. Figure 1). Note that the DAAO activity in the RPE1-hTert-DAAO^MLS^ and RPE1-hTert-DAAO^IMS^ was about half that of DAAO activity in RPE1-hTert-DAAO^H2B^ and RPE1-hTert-DAAO^TOM20^ cells at equal [D-Ala], which makes the difference in response even bigger.

**Figure 5:**
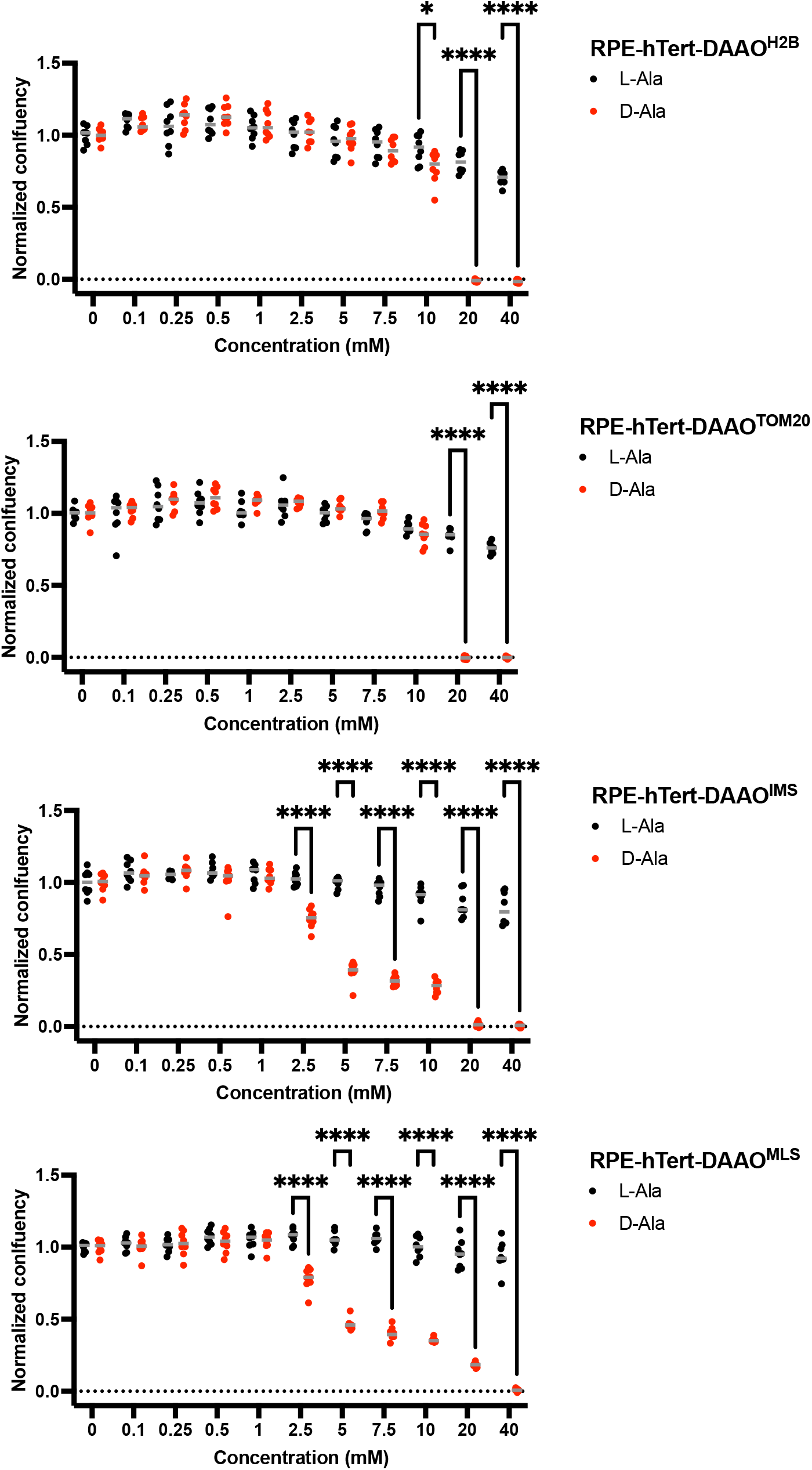
The subcellular site of H_2_O_2_ production determines its lethal dose Quantification of crystal violet assay shown in sup. figure 1 with RPE-1-hTert-DAAO lines treated with increasing concentrations of L- or D-Alanine for 48h. The decrease in confluency is not only dependent on the D-Ala concentration used, but also on the subcellular location of DAAO. Statistical significance was determined by 2-way Anova + Šídák’s multiple comparisons test (* = p < 0.05, **** = p < 0.0001, n = 8). A typical result of at least 3 replicate experiments with similar setup is shown.

## Discussion

The use of yeast D-amino acid oxidase has recently spurred research on intercompartmental H_2_O_2_ diffusion and signaling. However, as far as we know an assay that reports on the DAAO activity in live cells independently of the local reductive capacity has not been described before. The here described OCR-based DAAO activity assay can be used to compare DAAO activity across different cell lines, subcellular compartments, and D-alanine concentrations and gradients, irrespective of differences in local or global reductive capacity. This allows the establishment of DAAO model systems with comparable levels of H_2_O_2_ production, to distinguish phenotypic effects that stem from differences in for instance reductive capacity or total oxidative burden. By combining this assay with ultrasensitive, genetically encoded H_2_O_2_ sensors like roGFP2-Tsa2 and Hyper7, the (local) reductive capacity could be estimated by comparing the DAAO-dependent OCR with the Hyper7 signal, although it should be noted that these two parameters can as yet not be assessed simultaneously within the same well of cells. The use of oligomycin in the assay makes it possible to compare DAAO-dependent

H2O2 production to the amount of oxygen used in mitochondrial respiration. Mitochondrial respiration is a major source of ROS but the estimates of what percentage of oxygen used in mitochondrial respiration ‘escapes’ as superoxide and subsequently froms H_2_O_2_ are quite variable in literature. Often a value of 1-2% is reported based on pioneering work with isolated mitochondria (Beckman and Ames, 1998; Chance et al., 1979). More recently, however far lower estimations between 0.1% and 0.5% have been reported(Brand, 2016; Goncalves et al., 2015; Kudin et al., 2004; St-Pierre et al., 2002). Brand (Brand, 2016) even suggests that mitochondrial H_2_O_2_ generation is not directly related to the amount of mitochondrial respiration. It is therefore difficult to make a good estimate of what DAAO-dependent OCR is comparable to physiological levels of (mitochondrial) H2O2 production. However, at the levels of DAAO expression in our cell lines a substrate concentration of 2.5 mM D-Ala yields already more H2O2 than the maximum estimates mentioned in literature. At 10 mM D-Ala the amount of oxygen used for DAAO-dependent H2O2 production was as much as 7-20% of basal mitochondrial OCR. The exact percentage depended on the monoclonal line and may therefore not be directly translated to other systems using DAAO expression. Note that Oligomycin blocks ATP synthase and thereby inhibits OCR indirectly, whereas a combination of antimycin A and rotenone could potentially decrease mitochondrial OCR even further. We do however not expect that this would lead to much more accurate comparisons of DAAO-dependent OCR to mitochondrial OCR.

The OCR-based DAAO activity assay can in principle also be applied to other oxidases. Seahorse XF analyzers have previously been used to measure the activity of a genetically encoded NADPH oxidase, LbNOX (Cracan et al., 2017). Similarly, the activity of genetically engineered NADPH oxidase and H_2_O_2_ generator, P450 BM3, might also be estimated by measuring its oxygen consumption(Lim and Sikes, 2015). But since these enzymes use substrates that are, unlike D-amino acids, present normally in cultured cells it is more difficult to compare baseline and induced activity for these NADPH oxidases.

There are also some points to consider regarding the OCR-based DAAO activity assay. Firstly, it is difficult to measure levels of DAAO activity that are smaller than the variations in baseline OCR, even in oligomycin-treated cells. Increased oxygen consumption could reliably be measured upon addition of 5 mM D-alanine, but the corresponding levels of H_2_O_2_ production are probably already supraphysiological. At lower D-alanine concentrations H_2_O_2_ levels must therefore be extrapolated, which comes with the risk that for some reason the mathematical relationship between [D-Ala] and [H_2_O_2_] changes at [D-Ala] < 5 mM. Secondly, H_2_O_2_ that is reduced by catalase yields molecular oxygen, and therefore this fraction of H_2_O_2_ no longer contributes to OCR. The DAAO-dependent OCR may therefore underestimate the amount of H_2_O_2_ produced. Since catalase is localized to peroxisomes this may not be a major issue.

Besides these practical concerns, it is important to keep in mind that the OCR-based DAAO activity assay measures the amount of H_2_O_2_ produced by DAAO and not the resulting H_2_O_2_ concentration. If DAAO is localized to a relatively small subcellular compartment, a higher H_2_O_2_ concentration may be reached compared to equal DAAO activity in a larger compartment (assuming H_2_O_2_ does not diffuse out of the compartment).

Despite these limitations, we think that the method described here will contribute to a better understanding of DAAO-based model systems to generate H_2_O_2_ and by extension move the redox biology field forward.

## Materials

| REAGENT or RESOURCE | SOURCE | IDENTIFIER |
| --- | --- | --- |
| **Chemicals, peptides, and recombinant proteins** |  |  |
| 2-Deoxyglucose | Acros Organics | 111980250 |
| Blasticidin | Bio Connect | ant-bl-1 |
| D-alanine | Sigma-Aldrich | A7377 |
| Digitonin | Sigma-Aldrich | D141 |
| DMEM/F12 | Sigma-Aldrich | D8062 |
| DMEM High glucose | Sigma-Aldrich | D6429 |
| DMEM Low glucose | Sigma-Aldrich | D5546 |
| Fetal Bovine Serum | Bodinco | BDC-17748 |
| Glucose | Merck Millipore | 1.08337.1000 |
| L-alanine | Sigma-Aldrich | A7627 |
| L-glutamine | Lonza | 17-605E |
| NaOH | Merck Millipore | 1.06498.1000 |
| Oligomycin | Sigma-Aldrich | 75351 |
| Penicillin-streptomycin | Lonza | DE17602E |
| Polybrene | Sigma Aldrich | TR-1003 |
| Rat tail collagen 1 | Corning | 354236 |
| Saponin | Sigma-Aldrich | 47036-250G-F |
| Seahorse XF base medium | Agilent | 102353-100 |
| Seahorse XF Calibrant | Agilent | 102353-100 |
| Sodium Pyruvate | Sigma-Aldrich | S8636 |
| Trypsin | Sigma-Aldrich | T3924 |
| **Critical commercial assays** |  |  |
| Pierce™ BCA Protein assay kit | Thermo Scientific | 23225 |
| Seahorse XF24 V7 PS cell culture microplates | Agilent | 100777-004 |
| Seahorse XFe24 FluxPak | Agilent | 102340-100 |
| **Experimental models: Cell lines** |  |  |
| RPE1-hTERT H2B-DAAO |  |  |
| RPE1-hTERT IMS-DAAO |  |  |
| RPE1-hTERT MLS-DAAO |  |  |
| RPE1-hTERT TOM20-DAAO |  |  |
| RPE1-hTERT TOM20-DAAO + NES-HyPer7 |  |  |
| **Software and algorithms** |  |  |
| Fiji imaging software | https://fiji.sc/ |  |
| GraphPad Prism 9.3.1 | https://www.graphpad.com/scientific-software/prism/ |  |
| Wave Desktop and Controller 2.6 Software | https://www.agilent.com/en/product/cell-analysis/real-time-cell-metabolic-analysis/xf-software/seahorse-wave-desktop-software-740897 |  |
| **Other** |  |  |
| iMark Microplate Absorbance reader | Bio-Rad | 1681130 |
| Seahorse XFe24 analyzer | Aligent | 420017 |
| ZEISS Cell Observer microscope | ZEISS | N.A. |

## Methods

### Cell culture & Lentiviral transduction

RPE1-hTERT cells were cultured in DMEM/F-12 supplemented with 10% FBS, 2 mM L-glutamine and 100 Units Penicillin-Streptomycin. The cells were cultured at 37°C under a 6% CO2 atmosphere. The IDT (www.idtdna.com) codon optimization tool was used to convert the sequence DAAO from *Rhodotorula gracilis* for expression in human cells. The C-terminal peroxisomal targeting sequence was removed and a geneblock (see Supplementary information) was ordered for further cloning (Integrated DNA technologies). Fusions with the H2B-, TOM20-, IMS- and MLS-DAAO signals in a pLenti backbone were made by infusion cloning. DAAO expressing cells were generated using lentiviral infection and were subsequently made monoclonal by single-cell sorting. Generation of NES-HyPer7 lentiviruses and infection of TOM20-DAAO cells was performed as described previously (Shi et al., 2021). After infection, TOM20-DAAO + NES-HyPer7 cells with high HyPer7 fluorescence were single-cell sorted in conditioned medium and expanded to generate monoclonal cell lines.

### Determination of OCR using a Seahorse XFe analyzer

A Seahorse Bioscience XFe24 Analyzer was used to measure of oxygen consumption rates of RPE DAAO cells. A day before the assay, 24-well V7 Seahorse culture plates were coated with 50 μL of 50 μg/mL rat tail collagen 1 in 0.1% acetic acid for 20 minutes at room temperature. Afterwards the plate was washed with PBS and 40,000 cells were seeded in 100 μL medium. To 4 wells 100 μL of medium containing no cells was added to serve as background correction. After the cells had attached, an additional 150 μL of medium was added. In parallel, the Seahorse sensor cartridge was hydrated with Seahorse calibrant solution a day before the assay. Similarly, the Seahorse XFe24 analyzer was also turned on at this time to let it warm up overnight. On the day of the assay, cells were washed twice with assay media (XF base media supplemented with 2 mM L-glutamine, 17.5 mM glucose, 1 mM Sodium Pyruvate and 0.5 mM NaOH). Cells were incubated for 60 minutes in assay media in a non-CO_2,_ humidified incubator at 37 °C before starting the assay. Injections of L/D-alanine, 2 μM oligomycin, 25 μg/mL saponin and digitonin were used. The oxygen consumption rate was normalized to the oxygen consumption rate measured 25 minutes after oligomycin injection. For more details on the Seahorse-based DAAO activity assay see addendum 1.

### HyPer7 live imaging

For measuring HyPer7 oxidation by live microscopy, we refer to the method described previously by our laboratory (Soest et al., 2023).

### Crystal violet-based viability and outgrowth assay

Cells were counted after trypsinization and seeded in 96-well plates at a density of 0.5• 10^4^ cells per well. The next day, cells were exposed to a concentration range of [D-Ala] using the same concentrations of L-Ala as a control. D/L-Ala was left on the cells for 48 hrs. The plates contained 8 replicates for each cell line with a different DAAO localization and D/L concentration. Plates were washed twice with PBS, followed by fixation with icecold MeOH for 10 minutes. MeOH was aspirated and wells were overlaid with 0.5% w/v Crystal Violet in 25% MeOH for 10 minutes, followed by thorough washing with H_2_O. 8 empty wells per 96 well plate were used as blanks. Plates were air-dried overnight and scanned on an EPSON flatbed photo scanner. Crystal Violet was redissolved by adding 100 μL of 10% Acetic Acid per well and quantified by spectrophotometry (595 nm) using a Biorad iMark plate reader.

### Addendum 1: Step-by-step Seahorse DAAO activity assay protocol

In this step-by-step protocol, we will describe how to perform a Seahorse DAAO activity assay. This protocol is optimized for the XFe24 analyzer. However, in principle other Seahorse analyzers like the XF8 and XF96 analyzers can also be used. Similarly, while this protocol focuses on how to perform Seahorse DAAO activity assays on adherent cells, it can be adapted to non-adherent cells, organoids, *C. elegans* and even zebrafish embryos. Aligent maintains a searchable database of publications using the Seahorse. Using this database, recommended cell plate coatings, seeding densities, compound concentrations, etc. can easily be found.

### Day 1 Coating cell plate

Depending on the cell type, coating of the XF24 V7 PS cell culture plate may be necessary. Even cells that adhere well to regular cell culture plates, for example RPE1-hTERT cells, may require coating. Aligent provides cell culture plates pre-coated with Poly-D-Lysine. However, for RPE1 cells we have determined that coating with rat tail collagen 1 works best.

1. Prepare 1.3 mL of 50 μg/mL rat tail collagen 1 in 0.1% acetic acid.
2. Add 50 μL of the diluted collagen per well and incubate for at least 20 minutes at room temperature.
  a. More or less volume may be required in Seahorse plates with different surface areas.
3. Aspirate the collagen solution.
4. Wash the plate with PBS.
  a. Seed cells directly after aspirating the PBS.

### Seeding cells

The guideline for the number of cells to be seeded is that the basal respiration should be between 100 and 400 pmol/min. Furthermore, the maximal oxygen consumption (via respiration or DAAO activity) should not exceed 1000 pmol/min. The amount of cells to seed of course also depends on how much time the cells are allowed to grow. Typical seeding densities vary from 20,000 to 60,000 cells per well. For RPE1 cells it is best to seed 40,000 cells a day before the assay.

1. Harvest the cells.
2. Count the cells.
3. Dilute the cells to a concentration in which 100 μL medium contains the desired number of cells to seed.
4. Seed 100 μL of cell suspension per well. Do not seed cells in four of the wells. These wells will be used for background correction during the assay. Fill these wells with 100 μL of medium. For a homogenous cell monolayer, hold the pipette tip at an angle and rest the tip just below the circular rim on top of the well.
  a. Using this small amount of volume during seeding enhances the consistency of the monolayers. The cells, however, do require more medium to grow sufficiently overnight.
5. Place the plate in the incubator and allow the cells to adhere to the plate. This can take anywhere between 1 and 6 hours. RPE1 cells normally adhere in 2 hours.
6. Add 150 μL medium to all the wells when the cells have adhered. Be careful not to disturb the newly attached cells.
  a. Adding more than 150 μL of fresh medium is also possible if needed.

### Sensor cartridge and analyzer preparation

For correct functioning of the sensor cartridge, the probe tips must be hydrated in XF calibrant for 4 – 72 hours. A Seahorse FluxPak comes with a sensor cartridge, a utility plate to hydrate the probe tips and a Hydro booster to put between the sensor cartridge and utility plate.

1. Add 1 ml of XF calibrant to each well of the utility place and place the sensor cartridge with the Hydro booster on top of the utility plate. Ensure that all probe tips are submerged. Handle the sensor cartridge carefully since the probe tips are delicate.
2. Incubate the Sensor cartridge in a non-CO_2,_ humidified incubator at 37 °C for 4 – 72 hours. Humidification of the incubator prevents evaporation of the calibrant solution.
3. Turn on the Seahorse XF analyzer to allow it to equilibrate to the right temperature.

### Day 2 Preparing assay medium

Normally, Seahorse assays are performed in Seahorse XF Base Medium containing no bicarbonate or phenol red, since these compounds interfere with the pH measurements needed to quantify extracellular acidification. However, for the Seahorse DAAO activity assay only the oxygen consumption needs to be measured. It therefore may be possible to perform Seahorse DAAO activity assays in regular media. We have not tested this and therefore still recommend using Seahorse XF medium. Since Seahorse XF Base Medium also contains no glucose, glutamine, pyruvate or pH-buffer, supplementation is needed. For RPE cells we use Seahorse XF Base Medium supplemented with 2 mM L-glutamine, 17.5 mM glucose, 1 mM Sodium Pyruvate and 0.5 mM NaOH. To adapt the assay medium to another cell type, simply supplement Seahorse XF base medium to replicate the formulation of regular culture medium for that cell type. Avoid the use of serum, since this might interfere with oligomycin treatment and the injection ports.

1. Prepare 50 mL of assay medium and heat this to 37 °C. This is a sufficient volume for the washing of the cell plate and preparation of the injections. Due to the short assay time, the assay medium does not have to be prepared sterilely.

### Washing cell plate

To remove the old, regular medium from the cell plate, the cell plate has to be washed with assay media. Again, this might not be necessary for DAAO activity assays, but this has not yet been validated.

1. Aspirate all but 50 μL of medium.
2. Wash cells twice with 500 μL assay medium, leaving behind 50 μL of medium after each wash.
3. Add 475 μL of assay medium for a total of 525 μL of assay medium per well.
4. Observe the cells under the microscope to ensure that a uniform cell monolayer was maintained. Also make sure that all background correction wells remain empty.
5. Incubate the cells at 37 °C in a non-CO_2_, humidified incubator.

### Preparing injections

Per well, four different injection ports (A, B, C and D) on the sensor cartridge can be loaded. Injection ports can hold up to 100 μL, but the recommended volume is 75 μL. For injection, compounds are diluted in assay medium. It is important to note that each series of ports (For example all A ports) needs to be loaded with an equal amount of volume to ensure proper injection. For the DAAO activity assay, oligomycin can be given as the first injection. However, it is also possible to simply add oligomycin to the cell plate prior to the assay. This frees up an injection port. Instead of oligomycin, combined rotenone and antimycin A treatment can also be used to directly and more completely inhibit mitochondrial oxygen consumption. Several configurations are possible with L/D-amino acid injections. L-amino acid can be injected into all the wells as the first injection after oligomycin treatment or a selection of wells can be injected with L-amino acid while the rest are given D-amino acid. Similarly, multiple D-amino acid injections can be given consecutively to 1 well or to different wells in parallel. As an example, we will describe the loading of the injection ports with oligomycin, L-alanine and 2 D-alanine concentrations.

1. The first injection adds 75 μL to a volume of 525 μL (1/8). The desired concentration of oligomycin is 2 μM, we therefore load all A ports with 75 μL of 16 μM oligomycin.
2. The second injection adds 75 μL to a volume of 600 μL (1/9). The desired concentration of L-alanine is 10 mM, we therefore load all B ports with 75 μL of 90 mM L-alanine.
3. The third injection adds 75 μL to a volume of 675 μL (1/10). The desired concentration of D-alanine is 10 mM, we therefore load all C ports with 75 μL of 100 mM D-alanine.
4. The fourth injection adds 75 μL to a volume of 750 μL (1/11). The desired concentration of D-alanine is 20 mM. The well already contains 10 mM D-alanine. This will be diluted to 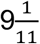 mM D-alanine by the 75 μL injection. To raise the D-alanine concentration to 20 mM, the concentration needs to be increased with 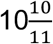 mM D-alanine. Load all D ports with 75 μL of 120 mM D-alanine.

### Design assay

Well, injection port and measuring configurations need to be inputted in Wave Desktop software before the assay. A guide for this can be found in section 2.4 of: https://www.agilent.com/en/product/cell-analysis/how-to-run-an-assay. One important thing to consider is the number of measurement cycles after an injection. Normally, three measurement cycles are performed after an injection, but it often takes longer for DAAO activity to stabilize. We therefore recommend six measurement cycles after each L/D-amino acid injection. Alternatively, the waiting time of measurement cycles can be increased.

### Run assay

1. Remove the Hydro booster and lid from the sensor cartridge. Put the sensor cartridge and calibrant-filled utility plate in the Seahorse Analyzer.
2. Wait roughly 15 minutes for the Seahorse Analyzer to calibrate.
3. When prompted, remove the utility plate from the Seahorse analyzer and place the cell plate (without the lid!) in the Seahorse analyzer. The assay will now start.

### Protein normalization

If protein normalization is preferred to normalization to oxygen consumption under oligomycin treatment, a BCA protein assay can be performed directly after the assay has been completed.

1. Wash the cell plate with PBS.
2. Lyse cells in 50 μL lysis buffer (0.1% Triton X-100, 10 mM Tris-HCl (pH 7.5), 5 mM EDTA and 100 mM NaCl in water).
3. Leave the plate in the fridge for at least an hour or in the freezer for overnight storage or longer.
4. Perform a protein assay according to the manufacturer’s protocol. We use a Pierce™ BCA Protein assay kit.

### Analysis

After the assay has been completed analysis can be performed in Seahorse Wave software. In this software, outliers can be removed, and oxygen consumption can be normalized to protein level or oxygen consumption under oligomycin treatment. The software will directly graph the results, but the data can also be exported to Excel or GraphPad Prism for further analysis.

