## Supplementary information for "Direct quantification of chemogenetic H_2_O_2_ production in live human cells"

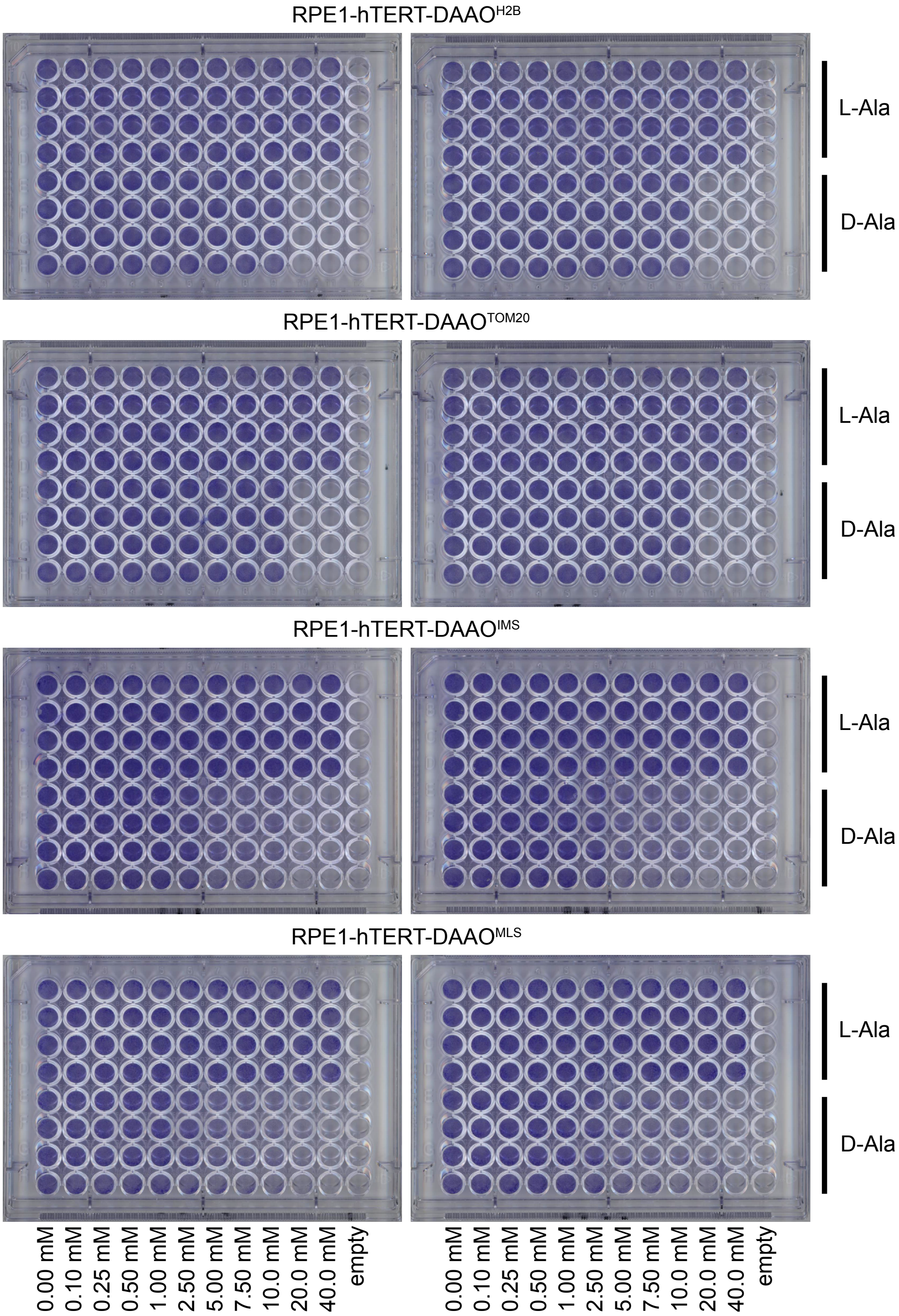


<u>Supplemental figure 1: Crystal violet assay with RPE1-hTert lines expressing differentially localized DAAO.</u>

*RPE1-hTert cell lines expressing differentially located DAAO were treated for 48h with several concentrations of L- or D-Ala (8 wells per condition in total). Quantification of this assay can be found in figure 5.*

Sequence of the used codon-optimized DAAO geneblock.

5’ACTCTCAAAAACGCGTCGTCGTTCTTGGCTCAGGAGTAATCGGGCTGTCATCCGCCC
TTATACTGGCCAGGAAGGGGTACTCAGTCCATATACTGGCAAGAGATCTCCCTGAAGA
CGTATCATCTCAAACGTTCGCCTCCCCCTGGGCTGGGGCGAACTGGACCCCCTTCATG
ACCCTGACAGATGGACCGCGGCAAGCAAAATGGGAGGAAAGTACGTTCAAGAAGTGG
GTCGAACTCGTCCCCACCGGCCACGCGATGTGGCTTAAAGGCACGCGCAGGTTTGCG
CAGAATGAGGACGGACTCCTTGGACATTGGTATAAAGATATAACTCCCAACTATCGACC
CTTGCCATCTTCAGAATGTCCCCCTGGCGCCATCGGTGTCACATACGATACGCTTAGTG
TGCACGCACCTAAATATTGCCAGTACCTCGCGCGGGAACTCCAAAAGTTGGGCGCAAC
GTTCGAGCGGAGAACAGTGACCTCTCTCGAACAAGCCTTCGATGGTGCGGACCTGGTT
GTTAACGCAACGGGCCTCGGGGCAAAGAGTATCGCTGGGATCGATGACCAAGCGGCT
GAACCAATACGCGGTCAGACAGTACTGGTAAAGTCACCCTGCAAAAGATGTACGATGG
ACAGTAGTGATCCAGCTTCCCCTGCTTATATTATTCCTCGACCAGGAGGTGAAGTTATA
TGTGGGGGGACTTATGGCGTGGGTGATTGGGACCTCTCAGTGAATCCAGAAACCGTGC
AGCGCATCCTGAAACACTGCCTGCGATTGGACCCAACAATAAGCTCAGATGGAACAATA
GAAGGAATTGAGGTCCTTAGACATAACGTCGGCCTCCGACCAGCTAGACGCGGAGGTC
CCAGGGTAGAGGCTGAAAGGATAGTTTTGCCTTTGGACCGCACCAAGAGTCCTTTGAG
TCTTGGACGCGGTTCTGCACGGGCAGCAAAAGAAAAGGAAGTTACCTTGGTTCATGCC
TACGGTTTCTCAAGCGCGGGGTATCAACAATCCTGGGGGGCGGCTGAAGATGTAGCCC
AACTTGTCGATGAGGCGTTTCAGAGATATCATGGGGCAGCCAGGGAGTAG-3’
